# Suicidal ideation and intentional self-harm in pregnancy as neglected agenda in maternal health; an experience from rural Sri Lanka

**DOI:** 10.1101/473058

**Authors:** Nimna Sachini Mallawa Archchi, Ranjan Ganegama, Abdul Wahib Fathima Husna, Delo Lashan Chandima, Nandana Hettigama, Jagath Premadasa, Jagath Herath, Harindra Ranaweera, Thilini Chanchala Agampodi, Suneth Buddhika Agampodi

## Abstract

**Background:** Suicide only present the tip of the iceberg of maternal mental health issues. Only a fraction of pregnant women with suicidal ideation proceeds to intentional self-harm (ISH) and even a smaller proportion are fatal. The purpose of the present study was to determine the prevalence of depression, suicidal ideation (present and past) and history of ISH among pregnant mothers in rural Sri Lanka.

**Methods:** We have conducted a hospital based cross sectional study in the third largest hospital in Sri Lanka and a another tertiary care center. Pregnant women admitted to hospital at term were included as study participants. The Edinburgh Postpartum Depression Scale (EPDS), a self-administered questionnaire for demographic and clinical data and a data extraction sheet to get pregnancy related data from the pregnancy record was used.

**Results:** The study sample consisted of 475 pregnant women in their third trimester. For the tenth question of EPDS “the thought of harming myself has occurred to me during last seven days” was answered as “yes quite a lot” by four (0.8%), “yes sometimes” by eleven (2.3%) and hardly ever by 13 (2.7%). Two additional pregnant women reported that they had suicidal ideation during the early part of the current pregnancy period though they are not having it now. Four (0.8%) pregnant women reported having a history of ISH during the current pregnancy. History of ISH prior to this pregnancy was reported by eight women and five of them were reported to hospitals, while others were managed at home. Of the 475 pregnant females included in the study, 126 (26.5%) had an EPDS score more than nine, showing probable anxiety and depression. Pregnant women who had primary/post-primary or tertiary education compared to those who were in-between those two categories were at higher risk of high EPDS with a OR of 1.94 (95% CI 1.1-3.3). Reported suicidal ideation prior to pregnancy had a OR of 6.4 (95% CI 2.3-17.5).

**Conclusions:** Based on our data, we conservatively estimate around 3000 ISH annually in Sri Lanka, which should be considered as a high priority for an urgent intervention.

**Plain English Summary:** Mental disturbances are common during pregnancy. Most of the time, these are normal. However, these disturbances may become serious and lead to self-harm and suicide. In this study, we estimated the proportion of pregnant women who had depression and idea of self-harming during pregnancy.

Respondents were pregnant women admitted to two large hospitals for the childbirth. They answered a list o questions about the thought of self-harm and attempts of self-harm during the present as well as past pregnancies.

Respondents included 475 pregnant women. Of them, 3.1% reported that “the thought of self-harming has occurred to them during last seven days quite a lot (0.8%) or sometimes” (2.3%). Four (0.8%) pregnant women reported that they actually did it to some extent. Of the 475 pregnant women included in the study, 126 (26.5%) had symptoms of anxiety/depression. Level of education seemed to have an association with anxiety and depression. When women reported that they had thought of self-harm prior to pregnancy, they were about 6.4 times more likely to have depression/anxiety during the pregnancy. Adding a simple screening question (as we used in this study) during the initial pregnancy assessment to detect history of suicidal thoughts will be helpful in identification of high-risk mothers for depression and suicide.

## Introduction

Ending preventable maternal mortality (EPMM) requires the consideration of all causes of maternal mortality, reproductive and maternal morbidities, and related disabilities[1][2]. Preventable direct obstetric causes are the main contributor to the global burden of maternal mortality (MM), whereas indirect causes account for 27.5% of maternal deaths and continuing to increase[3]. The concept of obstetric transition was proposed[4] and describes different stages in a dynamic process between and within countries. The transition (Stage I-V) ranges from high fertility and maternal mortality rates (MMR > 1,000/100,000 live births) to low fertility and MMR (<50/100,000 live births). It is also indicated that causes of MM shift in these stages from low access and quality of care, as well as direct obstetric causes to a high burden of indirect obstetric causes. While the global maternal health agenda offers a comprehensive approach towards EPMM[1], some components are still neglected in many settings.

Hitherto, there are no clear definitions of maternal suicide in many surveillance systems and the classification ranges from direct over indirect to incidental causes of maternal deaths between countries. The ICD-MM classification system clearly states that maternal suicides should be classified as direct maternal deaths[5]. Despite ICD-MM is in place since 2012, maternal suicides remain largely overlooked and inconsistently reported in many countries. A recent study in Sri Lanka found that over a period of six years, 17.8% of recorded maternal deaths in the North Central Province (NCP) were due to suicide, ranking it as the leading cause of MM[6]. This trend is also seen in other settings, especially in high income countries[7–9].

Suicide only present the tip of the iceberg of maternal mental health issues. Only a fraction of pregnant women with suicidal ideation precedes to intentional self-harm (ISH) and even a smaller proportion are fatal. However, suicidal ideation is shown to be one of the strongest predictors of suicide[10]. Contrary to the belief that pregnancy has a protective effect against suicide, a recent review clearly showed that pregnant women are more likely to harbor suicidal ideations[11]. The underlying maternal mental health issues leading to ISH and suicide are getting more attention due to increased knowledge on high prevalence of depression in pregnancy. The estimated prevalence of antenatal depression in low and middle income countries(LMIC) is 25.3% (21.4-29.6%)[12] and together with anemia, maternal depression is estimated to be the commonest cause of maternal morbidity[13]. In addition, prevalence of major depressive disorders in peripartum is estimated to be 11.9% with women from LMIC having a higher risk[14].

The primary focus of prevention strategies should be on improving maternal mental health, while the secondary prevention strategy, namely to reduce maternal suicides needs to concentrate on high-risk groups. Sri Lanka, considered a role model in reducing MMR, has yet to pay attention on reducing maternal suicide. Previous work on maternal depression clearly shows a high prevalence of antenatal[15,16] as well as postpartum depression[17,18] among Sri Lankan women. However, there is a lack of research on ISH or suicidal ideation among pregnant women in Sri Lanka. The purpose of the present study was to determine the prevalence of depression, suicidal ideation (present and past) and history of ISH among pregnant mothers in Anuradhapura as a mean to plan future strategies in EPMM in Sri Lanka.

## Methods

### Study settings

A hospital based descriptive cross-sectional study was carried out in the Teaching Hospital, Anuradhapura (THA) and the Base Hospital Thambuththegama (BHT). THA is the third largest hospital in the country with a bed capacity of more than 1,500. It covers a population size of more than 1.3 million and an average of 11,000 deliveries take place in this hospital annually. There are three obstetric units with five obstetricians (three from university unit and two from health ministry). BHT is situated 20km away from THA and can be considered as a main hospital, hosting a considerable number of deliveries. It has one obstetric unit functioning under an obstetrician. More than 80% of the deliveries in Anuradhapura district take place in these hospitals.

### Study sample

Study participants included all pregnant women with a gestational age more than 36 weeks (term pregnancies) admitted to antenatal wards of THA and BHT. This specific group was selected to maximize the period of exposure to collect the history of ISH. We excluded pregnant women who are illiterate, have disorders impairing their rational thinking, learning disabilities, are critically ill, and are in emergency by the time of admission.

### Sampling procedure

Consecutive women fulfilling inclusion criteria were included during the data collection period of a single month. Efforts were made to recruit mothers whose period of gestation could be calculated by confirmed dates by ultrasound scan. If this was not available, it was calculated from the start of amenorrhea.

### Variables

Main outcome variables were suicidal ideation (present and past) and history of intentional self-harm (during pregnancy and before pregnancy) as well as depression and anxiety.

### Data collection tools

The data collection instrument consisted of three main components. The Edinburgh Postpartum Depression Scale (EPDS) which is validated for the antenatal period, a self-administered questionnaire for demographic and clinical data and a data extraction sheet to get pregnancy related data from the pregnancy record which was used. The self-administered questionnaire was formulated in English and then translated to Sinhala and Tamil languages. An independent person who is fluent in both Sinhala and Tamil languages crosschecked the two questionnaires. Suicidal ideation was inquired using a single question “have you had thoughts of harming yourself “. This was asked for during pregnancy excluding the past week, previous pregnancies and before pregnancy. This was in addition to the EPDS question number 10.

### Data collection

Investigators visited all three obstetric wards in THA and the obstetric ward in BHT on a daily basis to recruit patients. The pregnant mothers who fulfilled the inclusion criteria were selected. They were provided with participation information leaflets. The objective of the study was explained, and they were given adequate time to ask questions regarding the study. After obtaining written informed consent, data collection was conducted by handing out the EPDS and the self-administered questionnaire s after giving clear instructions.

### Data analysis

Data were entered in an Epi-Info database. Data analysis was done using SPSS software. Prevalence and 95% confidence intervals for prevalence estimates were calculated. A binary logistic regression was used to predict depression.

### Ethical issues and patient concerns

If participants found to have symptoms of depression, all these patients were referred to the obstetrician for further assessment and referral, after a clinical interview. Ethical clearance for the study was obtained from the ERC committee of Faculty of Medicine and Allied Sciences, Rajarata University of Sri Lanka.

## Results

The study sample consisted of 475 pregnant women in their third trimester. The majority (n=444, 93.5%) was Sinhalese and from Anuradhapura district (n=440, 92.6%). The age of pregnant women ranged from 17 to 44 years with a mean of 28 (SD 6.5) years. There were 29 (6.1%) teenagers in the sample. Median family income was LKR. 30,000 per month (IQR 20,000-40,000).

History of miscarriage, intra uterine deaths (IUD) and stillbirths were reported by 75 (15.8%), four(.8%) and three(.6%) pregnant women. Medical illnesses or pregnancy complications during the current pregnancy were reported by 71 (14.9%) women (Table 2). In addition, ten participants reported family history of psychiatric illness and three reported family history of postpartum psychosis or depression.

**Table 1:** Characteristics of 475 pregnant women participated in the study.

|  | n | % |
| --- | --- | --- |
| **Age category** |  |  |
| <20 years | 34 | 7.2 |
| 20-34 years | 368 | 77.5 |
| >34 years | 73 | 15.4 |
| **Parity** |  |  |
| 1 | 172 | 38.8 |
| 2 | 180 | 40.6 |
| 3 | 73 | 16.5 |
| 4 | 13 | 2.9 |
| 5 | 5 | 1.1 |
| **Residence** |  |  |
| Anuradhapura | 440 | 92.6 |
| Kurunegala | 28 | 5.9 |
| Other areas | 7 | 1.4 |
| **Ethnicity** |  |  |
| Sinhalese | 444 | 93.5 |
| Muslim | 27 | 5.7 |
| Tamil | 4 | 1.4 |

| **Occupation** |  |  |
| --- | --- | --- |
| Housewife | 382 | 80.4 |
| Professionals | 50 | 10.5 |
| Semi professionals | 31 | 6.5 |
| Technical worker | 2 | 0.4 |
| Skilled laborer | 9 | 1.9 |
| Unskilled laborer | 1 | 0.2 |
| **Education Level** |  |  |
| Primary education not completed | 1 | 0.2 |
| Primary education completed | 38 | 8.0 |
| Post primary education completed | 230 | 48.4 |
| Secondary education completed | 155 | 32.6 |
| Diploma | 2 | 0.4 |
| Graduated | 49 | 10.3 |
| **Marital Status** |  |  |
| Married | 473 | 99.6 |
| Unmarried | 2 | 0.4 |

**Table 2:**
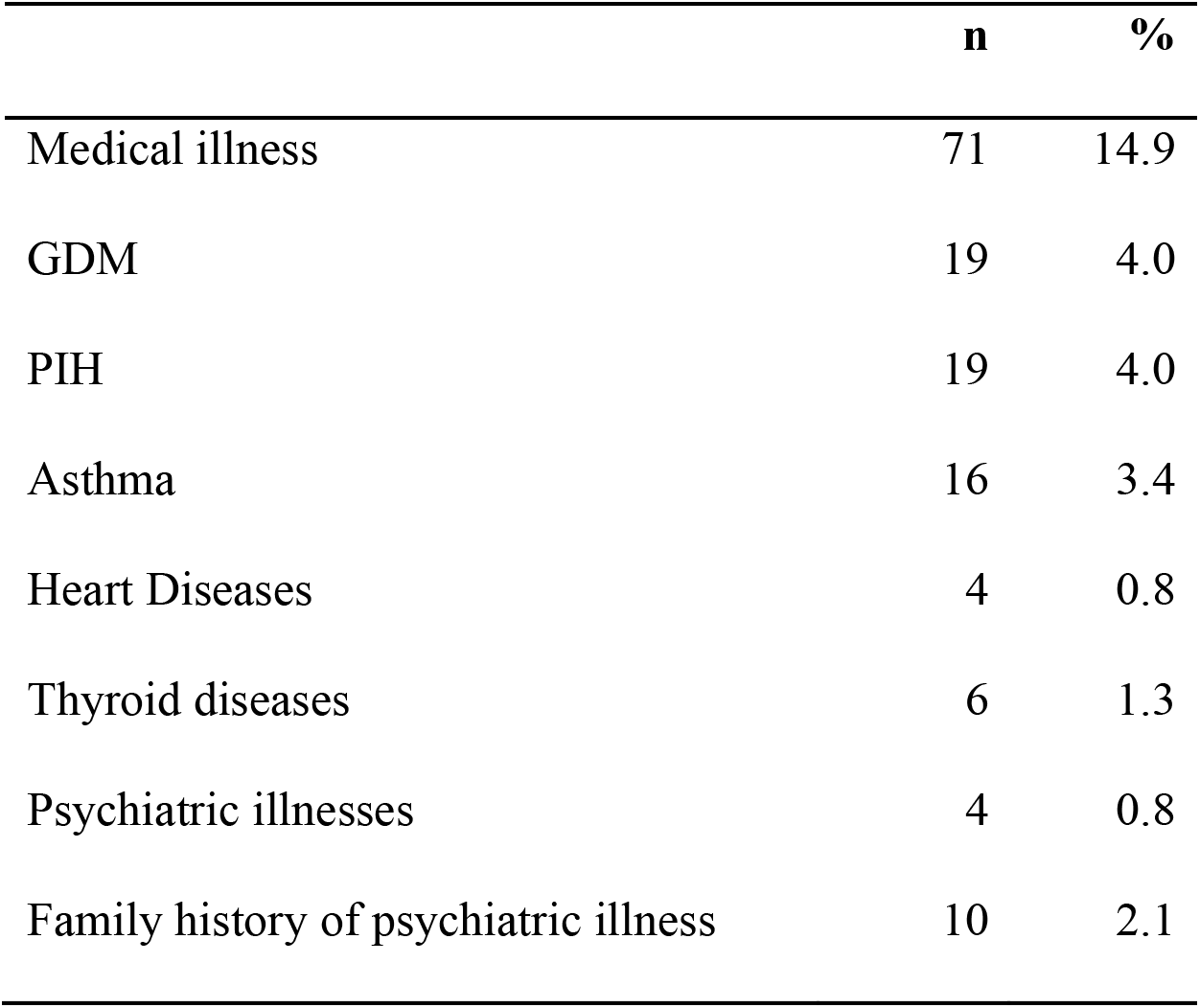
Self-reported medical conditions complicating pregnancy/pregnancy complications among 475 pregnant women in their third trimester.

**Table 3:** Estimates of self-reported suicidal ideation and intentional self-harm during the present pregnancy/lifetime.

|  | **Prevalence (%)** | **95% CI** |
| --- | --- | --- |
| Suicidal ideation in past seven days (EPDS) | 5.9 | 4.0 -8.4 |
| ISH during current pregnancy | 0.8 | 0.0 -1.7 |
| Suicidal ideation prior to current pregnancy | 3.8 | 2.1 -5.5 |
| ISH prior to current pregnancy | 1.7 | 0.5 -2.8 |

For the tenth question of EPDS “the thought of harming myself has occurred to me during last seven days” was answered as “yes quite a lot” by four (0.8%), “yes sometimes” by eleven (2.3%) and hardly ever by 13 (2.7%). Two additional pregnant women reported that they had suicidal ideation during the early part of the current pregnancy period though they are not having it now. Four (0.8%) pregnant women reported having a history of ISH during the current pregnancy (Table 2). All four of them were married, age ranges from 22 to 36 and two were primiparous. Of the two multiparous females with ISH during pregnancy, one had two ISH attempts; one during the previous pregnancy and another in-between pregnancies.

Three out of four (75%) who reported ISH during the current pregnancy had reported suicidal ideation even within the last seven days prior to the survey.

In three of the attempts agrochemicals were used, which are classified under ICD-10-CM Diagnosis Code T60.0X2A (Toxic effect of organophosphate and carbamate insecticides, intentional self-harm, initial encounter).

History of ISH prior to this pregnancy was reported by eight women, three (37.5%) had suicidal ideation during the current pregnancy. Two (25%) of them had ISH incidents during the current pregnancy (Figure 1). This was in comparison to 2/453 (0.4%) among those who were not having a history of ISH. Five out of eight (62.5%) previous ISH events were reported to hospitals, while others were managed at home.

**Figure 1:**
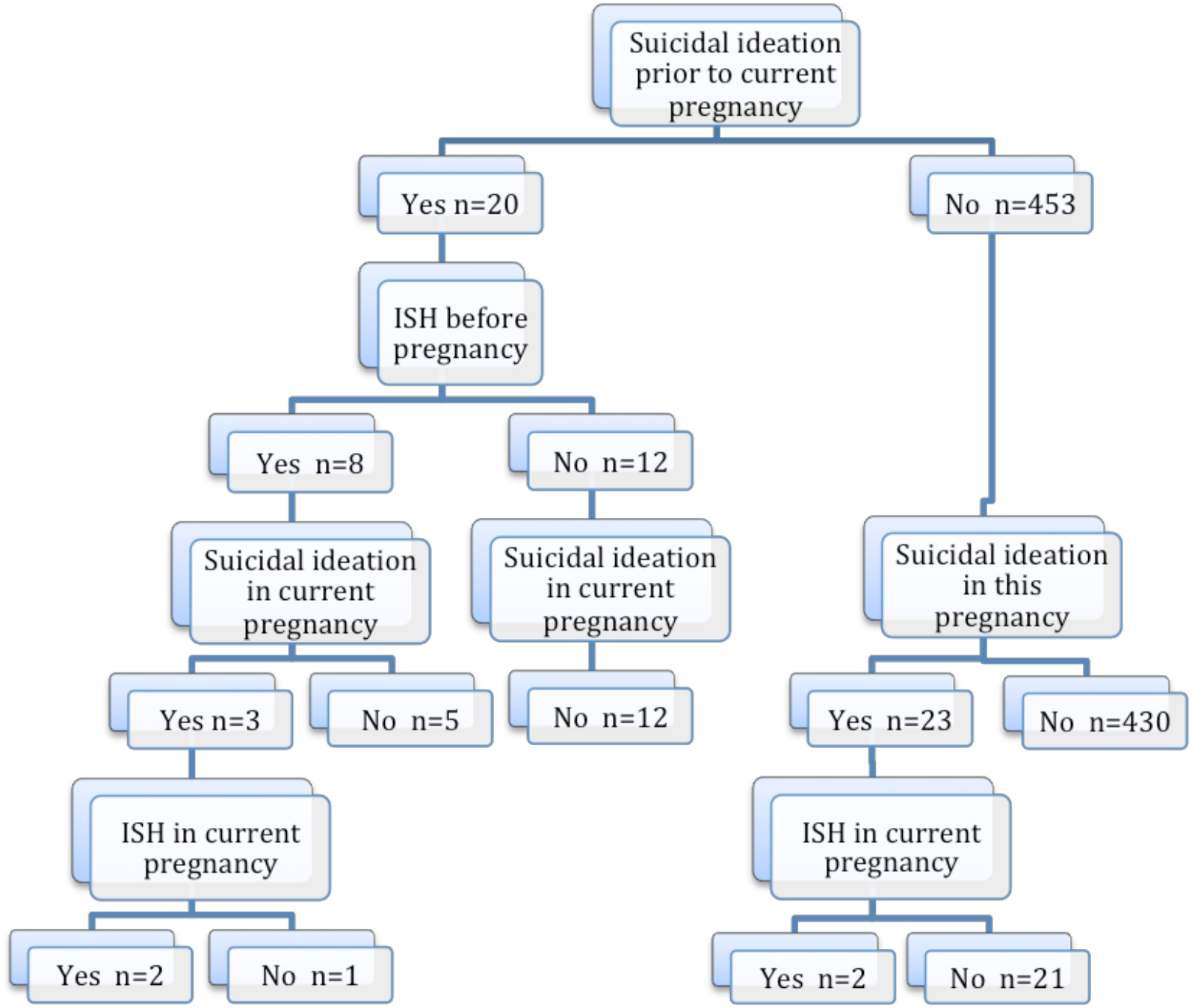
Suicidal ideation and intentional self-harm during the current pregnancy and prior to conception among 475 pregnant women at term.

Distribution of EPDS scores showed a median of six with interquartile range of four to ten. Of the 475 pregnant females included in the study, 126 (26.5%) had an EPDS score more than nine, showing probable anxiety and depression.

We looked at the risk factors to predict maternal depression and anxiety in this population using a binary logistic regression model. The explanatory variables used were age, parity, occupation, education, ISH prior to pregnancy and suicidal ideation prior to pregnancy. Pregnant women who had primary/post-primary or tertiary education compared to those who were in-between those two categories were at higher risk of high EPDS with a OR of 1.94 (95% CI 1.1-3.3). Reported suicidal ideation prior to pregnancy had a OR of 6.4 (95% CI 2.3-17.5).

## Discussion

The present study indicates a high burden of maternal mental ill-health and suicide risk among pregnant women in Sri Lanka, which is not reflected in routinely reported morbidity and mortality statistics. The last reported MMR in Sri Lanka was 33.8 per 100,000 live births[19] and the total count is around 121 maternal deaths. The number of maternal deaths attributed to suicide was less than 5% during 2005–13 period. However, applying findings of this study show that an estimated 5,000 pregnant women in Sri Lanka having suicidal ideations and around 3,000 commit ISH during pregnancy each year. Though the reported mortality due to underlying mental health issues is low, the actual burden of mental health issue in this study is estimated as very high.

Suicide among pregnant women is a preventable tragedy. Unlike the general population, pregnant women are under care of the health system and regularly examined by health professionals. Detection of suicidal ideation among pregnant women could be done without additional visits and minimal costs. Focusing on suicidal ideation, independently of depression, is important. It has been shown that suicidal ideation without clinical detectable mental health disorders are common[11]. Estimates of suicidal ideation show a wide range across countries and risk groups[20–23]. In Sri Lanka, among antenatal women in Anuradhapura it was reported as 6.9% in 2010[16] whereas during the postpartum period it was reported as 2.9%[18] (excluding “hardly ever”) in a larger sample from Sri Lanka, showing that the findings are consistent.

In this study, we assessed suicidal ideation during current pregnancy using a single question as well as a part of EPDS. It was noted that the single “yes/no” type question was grossly underestimating the past suicidal ideation. All pregnant women in the current study were in their third trimester. While 28 reported some degree of suicidal ideation within the last seven days, only six reported suicidal ideations prior to that. For previous pregnancies only one woman who had a ISH attempt during a past pregnancy reported suicidal ideation, among all 271 multiparous women. These numbers are not concordant and recall bias seemed to play a major role in this information. Any estimate beyond past two weeks of suicidal ideation or ISH may have resulted in an underestimation of the problem.

Even with probable underreporting, history of suicidal ideation is a strong predictor of high EPDS score as well as ISH. A simple screening question (as we used in this study) during the initial pregnancy assessment to detect history of suicidal ideation and ISH will be helpful in identification of high risk mothers for depression and suicide. Findings of this study need to be confirmed through a prospective study to capture ISH, as well as suicidal ideation throughout pregnancy after an initial assessment. Further, the sample size was small to predict a low prevalence, thus the role of chance cannot be excluded completely in these findings. The use of other tools for assessment of suicidal ideation could improve the quality of findings, however, the intention of this study focused on the feasibility of incorporating one single question as a risk assessment tool.

## Conclusion

The hidden burden on maternal mental health issues is a major concern for most of the LMIC in the region with similar socio-cultural background. The maternal health agenda in LMIC need a well-focused strategies to change the systems or programmes in order to tackle the issues related to maternal metal health. Adding screening tools or simple questions for risk identification needs to be considered as a priority.

## Declaration

### Ethics approval and consent to participate

All pregnant females provided informed written consent to participate in this study. Ethics approval was obtained from the Ethics Review Committee of Faculty of Medicine and Allied Sciences (ERC /2015 /17)

### Consent for publication

Not applicable

### Availability of data material

All data sets analyses during this study available from the corresponding author on reasonable request

### Authors’ contributions

SA perceived the study and helped in design, protocol writing, data analysis and interpretation. analyzed and interpreted data and wrote the manuscript. AWIPR, SMISJ, KMJ and SBU participated in planning, data collection and database preparation. SBA was involved in design, protocol preparation, data analysis and interpretation, and preparing the manuscript.

## Acknowledgments

The authors would like to acknowledge all pregnant female who volunteers for this study and the hospital staff of DGH Kagalle for their contribution to make the study a success and Ms. Constanze Friedl for editing the manuscript.

## Abbreviations

ISH: Intentional self-harm
EPDS: Edinburgh Postpartum Depression Scale
OR: Odds Ratio
CI: Confidence Interval
EPMM: Ending preventable maternal mortality
MM: Maternal Mortality
MMR: Maternal Mortality ratio
NCP: North Central Province
LMIC: Low and middle-income countries
THA: Teaching Hospital, Anuradhapura
BHT: Base Hospital Thambuththegama
LKR: Sri Lankan Rupees
IUD: Intra uterine deaths

